# Bacterial threat assessment of bacteriophage infection is mediated by intracellular polyamine accumulation and Gac/Rsm signaling

**DOI:** 10.1101/2022.04.01.486733

**Authors:** Camilla D. de Mattos, Artem A. Nemudryi, Dominick Faith, DeAnna C. Bublitz, Lauren Hammond, Margie A. Kinnersley, Caleb M. Schwartzkopf, Autumn J. Robinson, Alex Joyce, Lia A. Michaels, Robert S. Brzozowski, Alison Coluccio, Denghui David Xing, Jumpei Uchiyama, Laura K. Jennings, Prahathees Eswara, Blake Wiedenheft, Patrick R. Secor

## Abstract

When eukaryotic cells are killed by pathogenic microorganisms, damage-associated and pathogen-associated signals are generated that alert other cells of nearby danger. Bacteria can detect the death of their kin; however, how bacteria make threat assessments of cellular injury is largely unexplored. Here we show that polyamines released by lysed bacteria serve as damage-associated molecules in *Pseudomonas aeruginosa*. In response to exogenous polyamines, Gac/Rsm and cyclic-di-GMP signaling is activated and intracellular polyamine levels increase. In the absence of a threat, polyamines are catabolized, and intracellular polyamines return to basal levels, but cells infected by bacteriophage increase and maintain intracellular polyamine levels, which inhibits phage replication. Phage species not inhibited by polyamines did not trigger polyamine accumulation by *P. aeruginosa*, suggesting polyamine accumulation and metabolism are targets in the phage-host arms-race. Our results suggest that like eukaryotic cells, bacteria can differentiate damage-associated and pathogen-associated signals to make threat assessments of cellular injury.

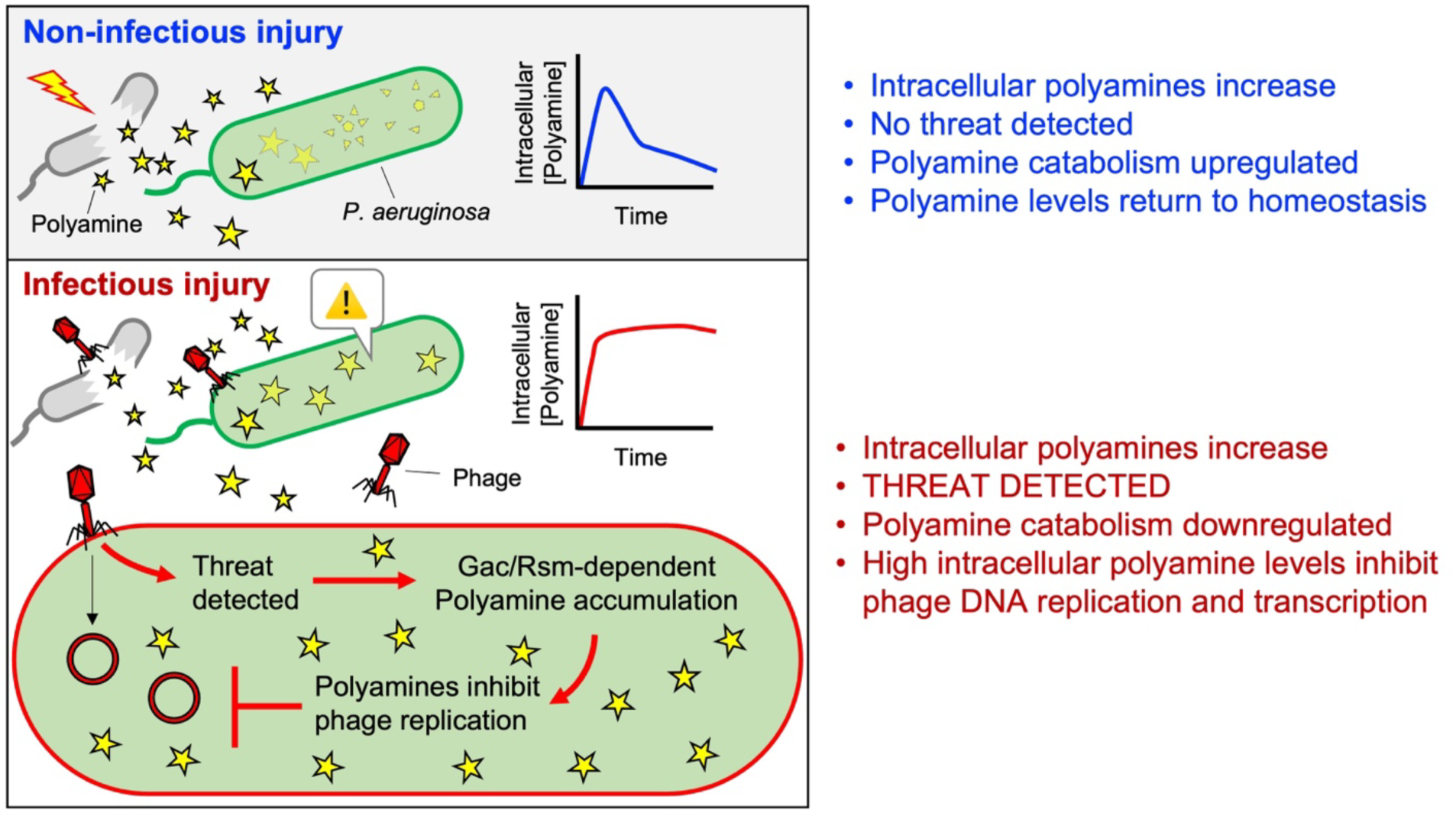

## Introduction

Bacteria have evolved innate and adaptive defense systems that protect them from viruses called bacteriophages (phages). Examples of phage defense systems include restriction modification (Kruger and Bickle, 1983), CRISPR-Cas systems (Wiedenheft et al., 2012), and abortive infection systems that kill or inhibit the bacterial host before the phage can complete its lifecycle (Bernheim and Sorek, 2019; Cohen et al., 2019; Doron et al., 2018). Recent comparative immunological studies reveal some bacterial abortive infection systems share evolutionary roots with eukaryotic antiviral immune pathways (Cohen *et al*., 2019; Ofir et al., 2021). Bacterial and eukaryotic immune systems share other conceptually analogous characteristics, such as the ability to detect molecules that are indicative of cellular damage.

When eukaryotic cells lyse, molecules such as ATP, intracellular proteins, oxidized lipids, and others, are released and sensed by conserved pattern recognition receptors (PRRs) on neighboring cells (Matzinger, 2002; Zhivaki and Kagan, 2021). PRRs also detect microbial-derived molecules and stimulate antimicrobial immune responses (Janeway, 1989). Eukaryotic cells take cues from both damage- and microbial-derived molecules to make threat assessments of cellular injury (Zhivaki and Kagan, 2021). Bacteria also sense and respond to molecules released by lysed bacteria (Bhattacharyya et al., 2020; LeRoux et al., 2015a; LeRoux et al., 2015b; Tzipilevich et al., 2021). However, the molecules that serve as damage signals in bacteria are poorly characterized and how they influence phage defense remains largely unexplored.

Here, using the human pathogen *Pseudomonas aeruginosa* as a model system, we identify a new phage defense system in which polyamines released by lysed bacteria function as a danger signal preventing phage-mediated cell death in neighboring bacteria. Polyamines are internalized by surviving cells, causing intracellular polyamine levels to increase, a process that is dependent on Gac/Rsm and cyclic-di-GMP signaling. In the absence of a threat of phage infection, intracellular polyamines are catabolized and return to basal levels. However, when bacteria are infected with phage, intracellular polyamine levels remain elevated, which inhibits phage genome replication and transcription. Collectively, our study supports a model where bacteria exploit polyamines to sense cell damage and inhibit phage replication.

## Results

### A small water-soluble molecule released by lysed bacteria suppresses phage replication

Phage infections can result in mass bacterial cell lysis. We hypothesized that lysed bacteria would release a soluble signal that would induce phage defense mechanisms in neighboring cells. To test this hypothesis, we pelleted and washed *P. aeruginosa* cells, resuspended them in fresh LB broth, and lysed them by sonication. After removing cell debris by centrifugation, the crude soluble lysate was used to make LB agar plates. Lawns of *P. aeruginosa* PAO1 grown on LB agar or LB-lysate agar were then challenged with phage species representing families Siphoviridae (JBD26, DMS3vir), Podoviridae (CMS1), Myoviridae (F8), and Inoviridae (Pf4). Phage titers were measured by quantifying plaque forming units (PFUs) on lawns of bacteria. Replication of all phages except CMS1 were inhibited on LB-lysate agar (**Fig 1A**). When *P. aeruginosa* collected from LB-lysate agar plates were re-plated onto LB agar, sensitivity to phage infection was restored (**Fig 1B**), indicating that cell lysates induce a transient phage tolerance phenotype in *P. aeruginosa* rather than heritable mutations that confer phage resistance.

**Figure 1.**
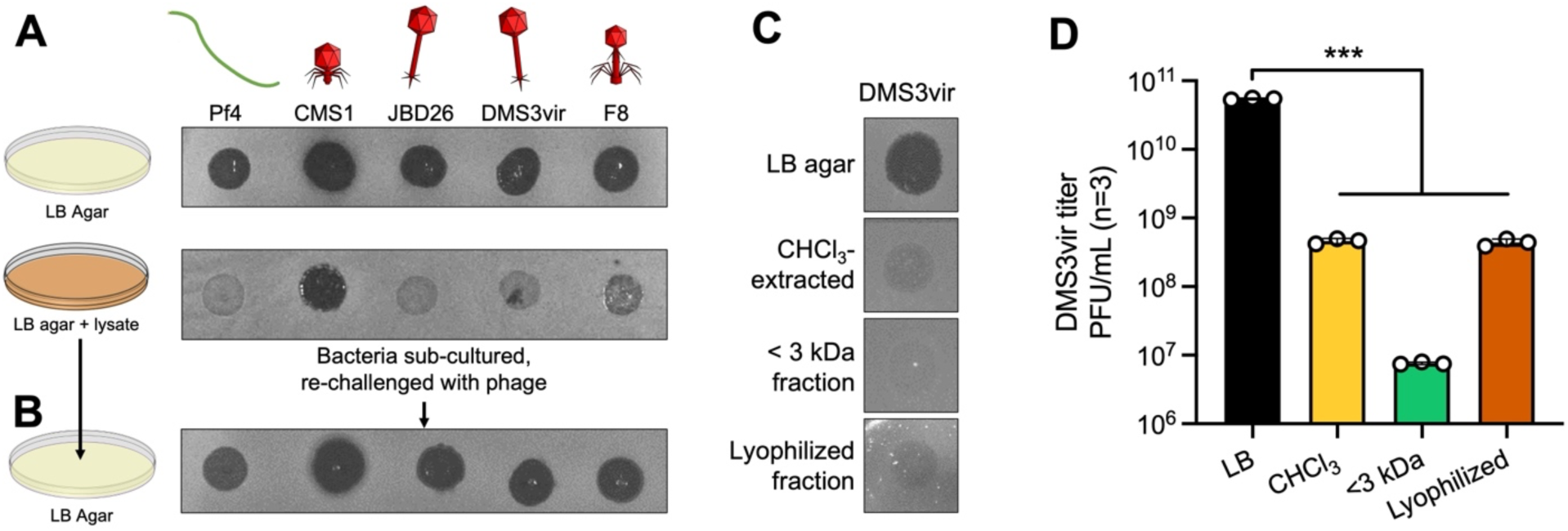
Small soluble molecules released by lysed *P. aeruginosa* cells induce transient tolerance to phage infection. **(A)** The indicated phage species were spotted at 10^6^ PFUs in 3 µl onto lawns of *P. aeruginosa* PAO1 grown on LB agar or LB agar supplemented with filtered PAO1 cell lysate. Plaques were imaged after overnight (18 h) growth at 37ºC. Representative images are shown. **(B)** *P. aeruginosa* growing on lysate plates were then sub-cultured onto LB agar plates and re-challenged with the indicated phages. Representative images are shown. **(C)** Phage DMS3vir was spotted at 10^6^ PFU in 3 µl onto lawns of *P. aeruginosa* growing on LB agar or LB agar supplemented with the aqueous phase of CHCl_3_-extracted cell lysate, the aqueous phase passed through a 3 kDa membrane, or lyophilized aqueous phase extract. Representative plaque images are shown. **(D)** DMS3vir titers were enumerated by plaque assay. Results are mean ± SD of three experiments, ***P<0.001.

To determine if the molecule(s) responsible for inducing phage tolerance are hydrophobic or water-soluble, we extracted LB lysate with chloroform (CHCl_3_) and used the aqueous phase to make agar plates. We repeated the plaque assays using DMS3vir, a lytic mutant of the *Mu*-like temperate *Pseudomonas* phage DMS3 (Cady et al., 2012). The active molecule(s) were retained in the aqueous phase, were able to pass through a 3 kDa molecular weight cutoff membrane, and retained activity after lyophilization and rehydration (**Fig 1C and D**). Together, these results indicate that small water-soluble molecule(s) are inducing a phage tolerance phenotype in *P. aeruginosa*.

### The Gac/Rsm pathway is required for lysate-induced phage tolerance in *P. aeruginosa*

Prior work demonstrates that a soluble signal released by lysed *P. aeruginosa* is sensed by the Gac/Rsm pathway in kin cells (LeRoux *et al*., 2015a). In *P. aeruginosa*, Gac/Rsm regulates many bacterial behaviors related to biofilm formation and virulence at the transcriptional level (Ventre et al., 2006) (**Fig 2A**). Gac/Rsm signaling is initiated by the activation of the sensor histidine kinase GacS by unknown ligands (Latour, 2020). GacS phosphorylates the response regulator GacA, which induces the transcription of the small RNAs *rsmY* and *rsmZ* (Brencic et al., 2009). These sRNAs bind and sequester the mRNA-binding proteins RsmA or RsmN away from their target mRNAs, allowing the translation of hundreds of different mRNAs (Lapouge et al., 2008; Romero et al., 2018). The sensor kinase RetS counteracts GacS activity and deletion of the *retS* gene constitutively activates Gac/Rsm signaling (Francis et al., 2018; Goodman et al., 2009; Mougous et al., 2006).

**Figure 2.**
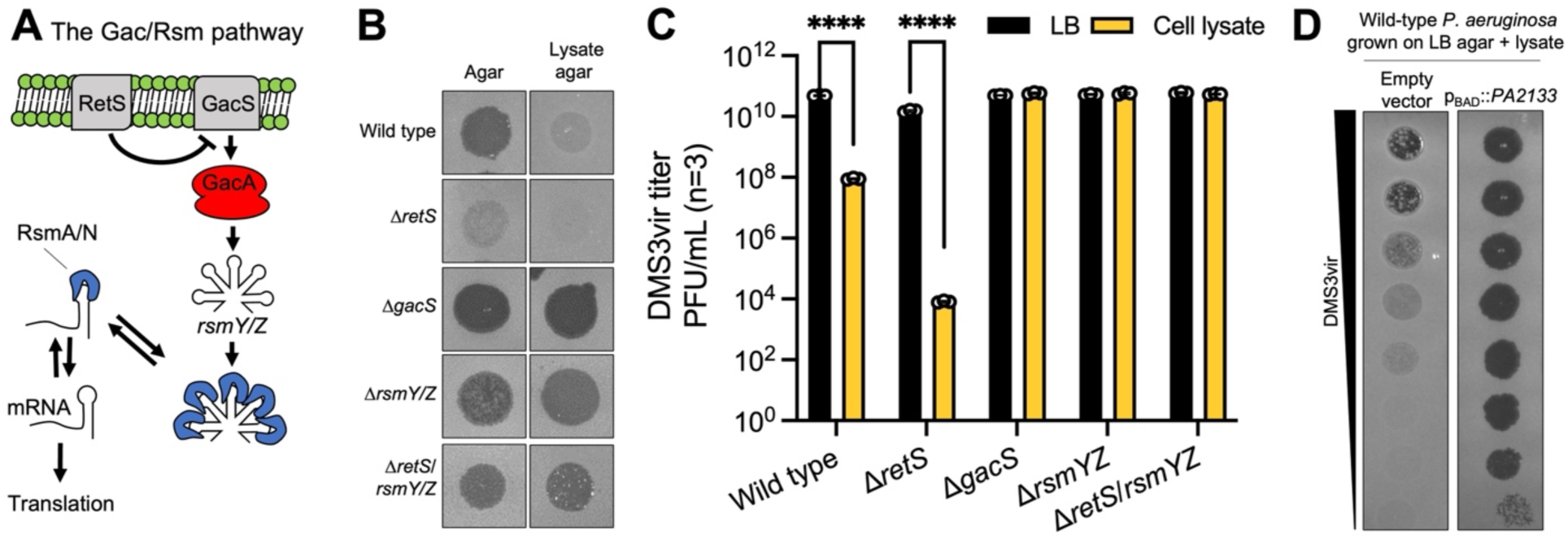
Phage tolerance is dependent upon Gac/Rsm and cyclic-di-GMP signaling. **(A)** Schematic of the Gac/Rsm pathway in *P. aeruginosa*. **(B and C)** Phage DMS3vir was spotted at 10^6^ PFU in 3 µl onto lawns of the indicated strains grown on LB agar or LB lysate agar. Representative plaque images are shown in (B) and PFU measurements are shown in (C). Results are mean ± SD of three experiments, ****P<0.0001. **(D)** Serial dilution plaque assays comparing the plating efficiency of phage DMS3vir on *P. aeruginosa* carrying an inducible c-di-GMP-degrading phosphodiesterase (p_BAD_::*PA2133*) or a control strain carrying an empty vector. Bacterial lawns were grown on LB agar supplemented with CHCl_3_-extracted lysate and 0.1% arabinose for 18 h.

We hypothesized that the Gac/Rsm pathway would be essential for cell lysate to induce phage tolerance in *P. aeruginosa*. To test this hypothesis, we used strains of *P. aeruginosa* where the Gac/Rsm pathway was either disabled (Δ*gacS,ΔrsmY/Z*) or constitutively activated (Δ*retS*). On LB agar, phage DMS3vir formed clear plaques on wild type, Δ*gacS*, and Δ*rsmY/Z* lawns, but formed turbid plaques on Δ*retS* lawns (**Fig 2B**, left column), suggesting that active Gac/Rsm signaling in the Δ*retS* strain is sufficient to impede DMS3vir replication. On LB-lysate agar, DMS3vir infection was suppressed on wild-type lawns and completely inhibited on Δ*retS* lawns (**Fig 2B and C**); however, sensitivity to DMS3vir infection was restored in strains where Gac/Rsm signaling was disabled (Δ*gacS*, Δ*rsmY/Z*) (**Fig 2B and C**). Furthermore, deleting *rsmY* and *rsmZ* from the phage-tolerant Δ*retS* background (Δ*retS/rsmY/Z*) restored susceptibility to phage infection on LB-lysate agar (**Fig 2B and C**). These results indicate that the Gac/Rsm pathway is essential for lysate-induced phage tolerance.

In *P. aeruginosa*, activation of Gac/Rsm signaling increases intracellular levels of the second messenger cyclic-di-GMP (Moscoso et al., 2011). In *P. aeruginosa*, cyclic-di-GMP regulates processes involved in the transition between motile (low cyclic-di-GMP) and sessile (high cyclic-di-GMP) lifestyles (Valentini and Filloux, 2016). To test the hypothesis that cyclic-di-GMP signaling is required for lysate-induced phage tolerance, we expressed the phosphodiesterase PA2133, which rapidly degrades cyclic-di-GMP in *P. aeruginosa* (Hickman et al., 2005). Expressing PA2133 in wild-type *P. aeruginosa* on lysate agar plates restored sensitivity to DMS3vir (**Fig 2D**). These results provide a link between cyclic-di-GMP signaling and phage defense in *P. aeruginosa*.

### Polyamines induce Gac/Rsm-dependent phage tolerance

We used RNA-seq to determine how CHCl3-extracted lysate affected the transcriptional profile of *P. aeruginosa* (**Supplemental Table S1**). Compared to cells growing on LB agar, 394 genes were differentially regulated in *P. aeruginosa* growing on LB-lysate agar (**Fig 3A**). Cell lysate also upregulated the small RNAs *rsmY* and *rsmZ* (**Fig S1**), indicating that cell lysate activates Gac/Rsm signaling. Gene enrichment analysis revealed upregulated genes associated with spermidine/polyamine catabolism (breakdown) are over-represented in this dataset (**Fig 3B**), several of which are highlighted in the volcano plot shown in Figure 3A.

**Figure 3.**
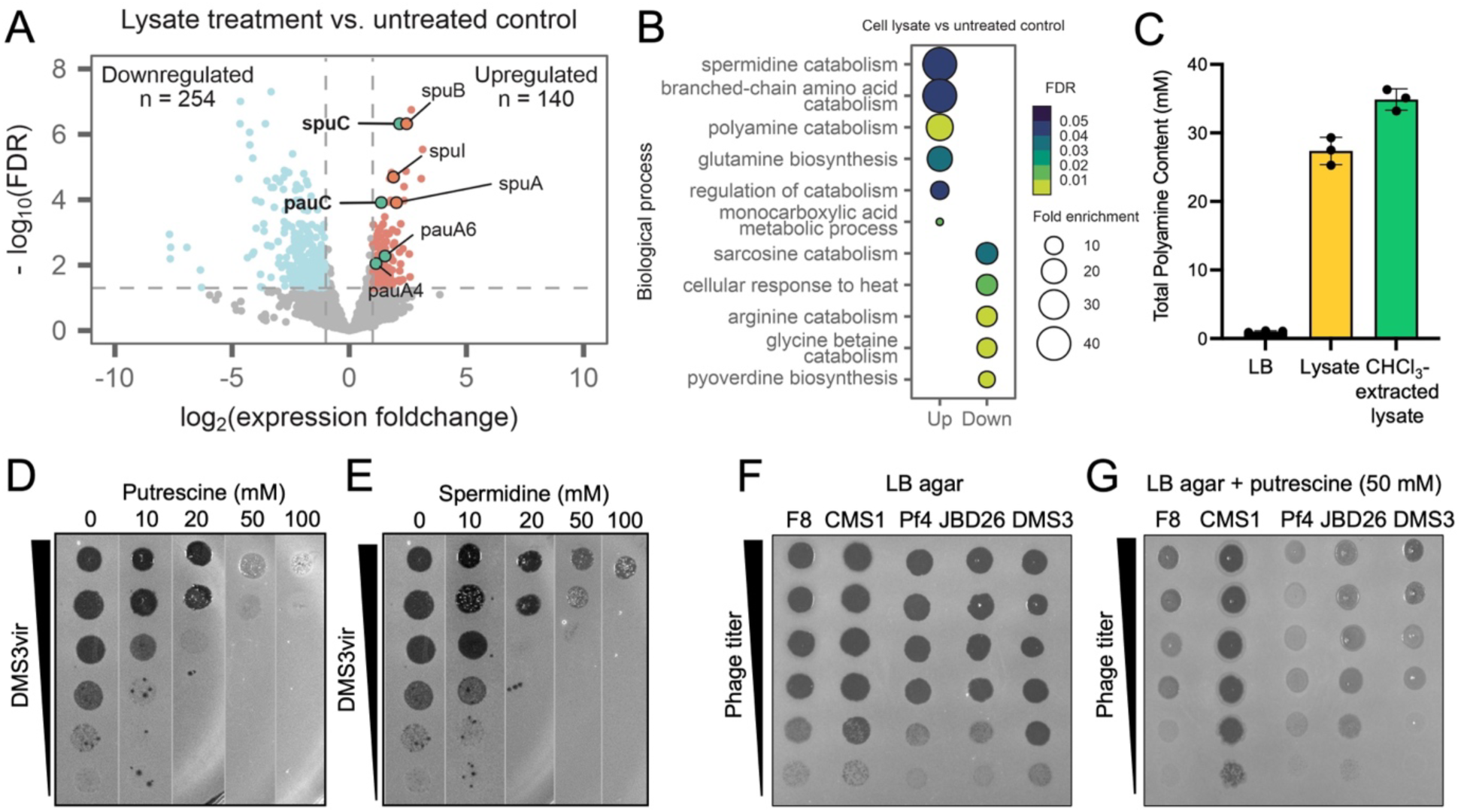
The polyamines putrescine and spermidine induce phage tolerance in *P. aeruginosa*. **(A)** Volcano plot showing differentially expressed genes in cells treated with lysate compared to untreated controls. Red indicates upregulated genes (log_2_[foldchange] > 1 and FDR<0.05) and blue indicates downregulated genes (log_2_[foldchange] > 1 and FDR<0.05). Non-significant genes are shown in gray. (**B)** Gene enrichment analysis of significant differentially expressed genes shown in (A). Dot sizes indicate fold enrichment of observed genes associated with specific Gene Ontology (GO) terms verses what is expected by random chance. **(C)** Total polyamine content in LB broth, cell lysate, and CHCl_3_-extracted cell lysate were measured using a fluorometric assay. Results are mean ± SD of three experiments, **P<0.01. **(D and E)** The polyamines putrescine (D) or spermidine (E) were added to LB agar at the indicated concentrations. DMS3vir was spotted onto lawns of *P. aeruginosa*, and plaques were imaged after 18 h of growth at 37ºC. (**F and G**) The indicated species of phage were spotted onto lawns of *P. aeruginosa* growing on LB agar or LB agar supplemented with 50 mM putrescine.

**Figure S1.**
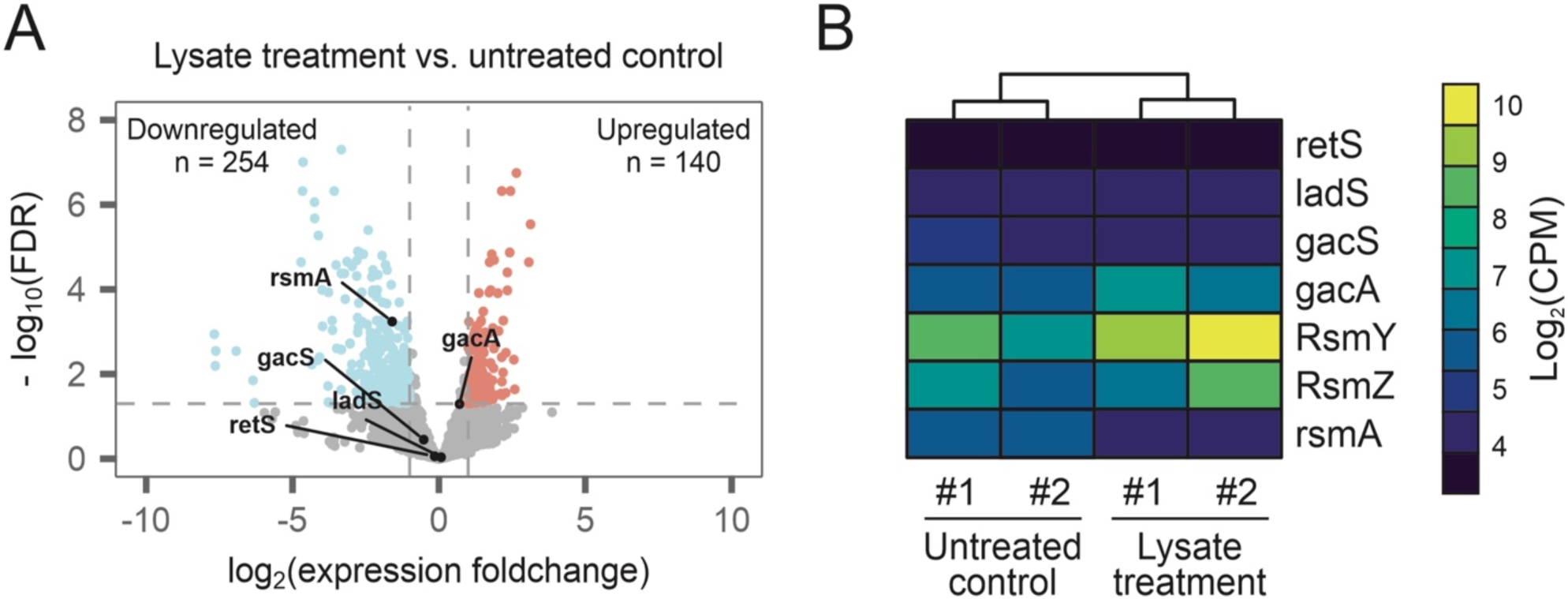
Cell lysate exposure upregulates *rsmY* and *rsmZ* regulatory RNAs. **(A)** Volcano plot showing DEGs in cells treated with lysate. Gac/Rsm pathway protein coding genes are indicated. **(B)** Heatmap showing expression of Gac/Rsm pathway members. Coloring indicates log_2_ of counts per million (CPM) reads. Two biological replicates for each treatment are shown.

Polyamines are small polycationic compounds that are involved in a variety of cellular processes in eukaryotes and prokaryotes through interactions with negatively charged molecules such as DNA, RNA, and proteins (Childs et al., 2003; Sarkar et al., 1995). In bacteria, putrescine and spermidine are the most common polyamines (Banerji et al., 2021) and are typically present at high concentrations (0.1–30 millimolar) inside bacterial cells (Duprey and Groisman, 2020; Shah and Swiatlo, 2008). In response to phage infection or other mass-lysis events, millimolar concentrations of polyamine could be released into the immediate environment. Indeed, in the cell lysates we used to make LB-lysate agar plates, total polyamine concentrations were approximately 30 mM (**Fig 3C**).

To test the hypothesis that polyamines would inhibit phage replication, we grew *P. aeruginosa* lawns on LB agar (MOPS buffered at pH 7.2) supplemented with 0, 10-, 20-, 50-, or 100-mM putrescine or spermidine. Both polyamines suppressed DMS3vir replication in a dose-dependent manner (**Fig 3D and E**). Putrescine also inhibited replication of phages Pf4, JBD26, and F8, but not CMS1 (**Fig 3F and G**), consistent with our observations using crude bacterial lysates (**Fig 1A**). Exposure of *P. aeruginosa* to 50 mM putrescine upregulated polyamine catabolism genes (**Fig S2**), also consistent with experiments performed using cell lysate.

**Figure S2.**
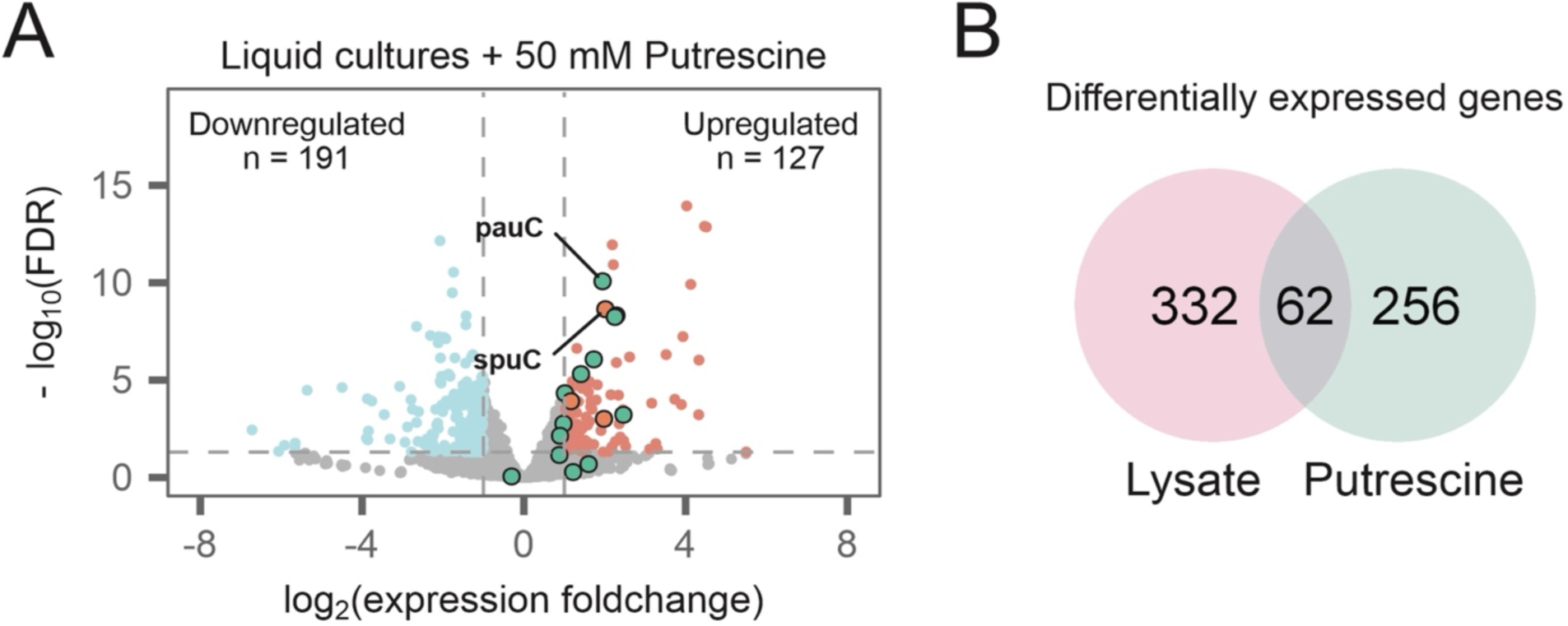
Putrescine activates polyamine catabolic pathways. **(A)** Volcano plot of RNA-seq data showing differentially expressed genes in wild-type *P. aeruginosa* cultured in liquid LB cultures with addition of 50 mM putrescine. Red dots indicate upregulated genes (log_2_(foldchange) >1 and FDR<0.05), and blue indicates downregulated genes (log_2_(fold change) <-1 and FDR<0.05). Non-significant genes are shown in gray. Genes involved in polyamine and putrescine catabolism are highlighted. **(B)** Venn diagram showing differentially expressed genes overlapping between putrescine and cell lysate treatments.

When grown in LB broth supplemented with 50 mM putrescine, wild-type and Δ*retS P. aeruginosa* were less susceptible to DMS3vir infection (**Fig 4A and B**, dashed lines) while disabling Gac/Rsm signaling (Δ*gacS*) restored sensitivity to DMS3vir infection (**Fig 4C**, dashed line). Upon visualization of cells, we noted that wild-type and Δ*gacS* cells infected by DMS3vir were on average significantly longer (2.6 µm) than uninfected cells (1.9 µm) (**Fig 4D and E**). Supplementing exogenous putrescine prevented DMS3vir infection-induced cell elongation. In the Δ*retS* strain, which is inherently tolerant to phage infection, cell length did not significantly change under any condition tested (**Fig 3D and E**). These results suggest that successful infection by DMS3vir inhibits *P. aeruginosa* cell division, but not growth, resulting in cell elongation. These results also suggest that putrescine disrupts DMS3vir replication before the phage can suppress *P. aeruginosa* cell division—so long as Gac/Rsm signaling is intact.

**Figure 4.**
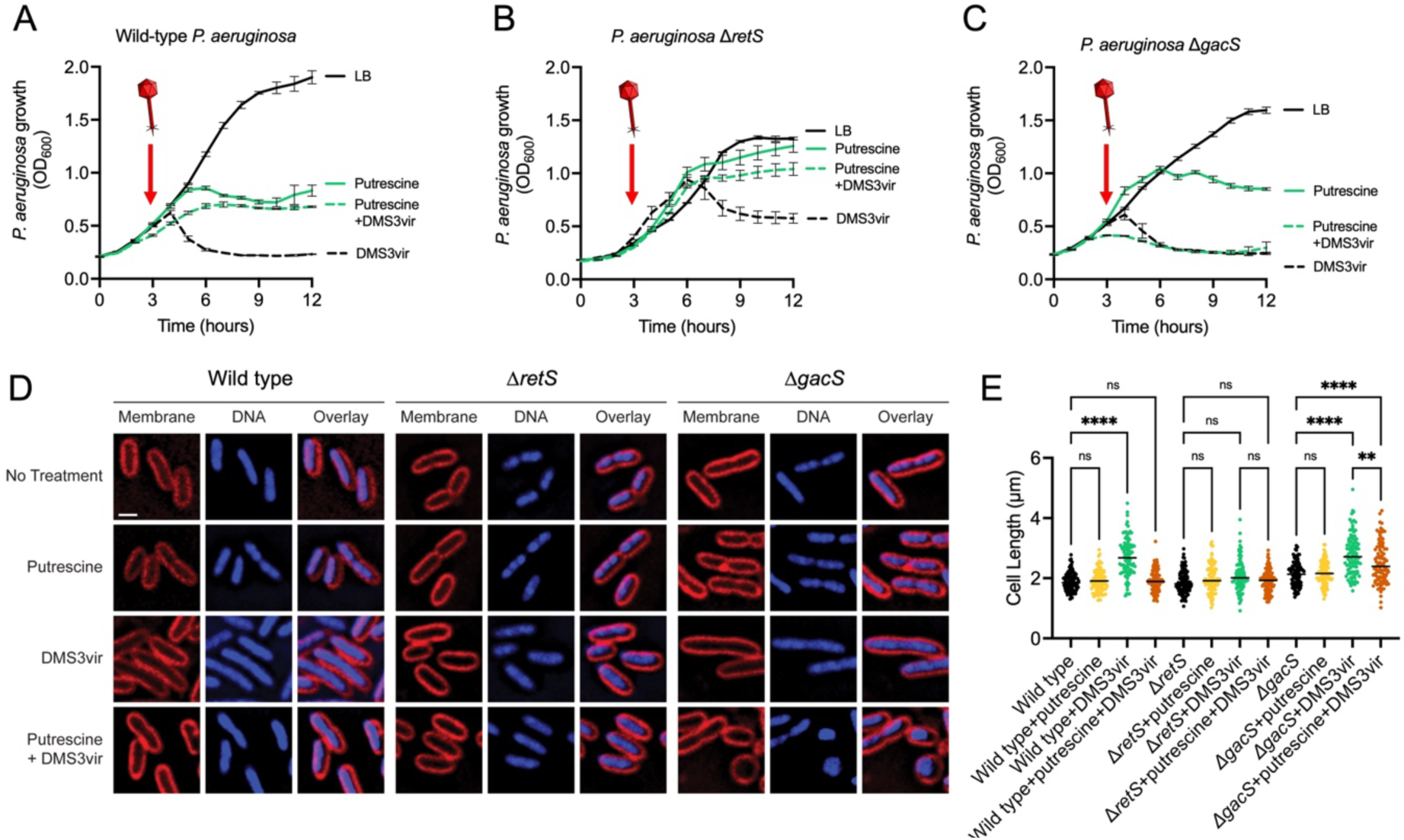
Putrescine induces Gac/Rsm-dependent tolerance to DMS3vir infection. **(A-C)** Growth curves of the indicated *P. aeruginosa* strain in LB broth or LB supplemented with 50 mM putrescine with and without infection by phage DMS3vir (arrow) at a multiplicity of infection (MOI) of 1. For each experiment, two biological replicates are presented as individual curves, each a mean of two technical replicates. (**D**) Fluorescence micrographs of wild-type, Δ*retS*, or Δ*gacS P. aeruginosa* strains stained with SynaptoRed (membrane; red) and DAPI (DNA; blue) 2 h post-treatment with putrescine, phage DMS3vir, both, or neither. Scale bar is 1 µm. (**E**) Quantification of cell length of cells in each treatment condition shown in panel D. n = 100 cells; **** P < 0.0001, ** P = 0.0053, and ns = not statistically significant.

Putrescine suppressed the growth of wild-type *P. aeruginosa* compared to bacteria growing in LB (**Fig 4A**, solid lines). It is possible that suppressed bacterial growth by polyamines explains phage tolerance. However, Δ*gacS* also displayed polyamine-induced growth suppression but remained sensitive to phage infection (**Fig 4C**, solid lines), indicating that growth inhibition by putrescine alone does not promote tolerance to DMS3vir infection. Furthermore, expression of the phosphodiesterase PA2133 to degrade intracellular cyclic-di-GMP levels re-sensitized wild-type and Δ*retS P. aeruginosa* to DMS3vir infection in the presence of 50 mM putrescine (**Fig S3**). Collectively, these results indicate that Gac/Rsm and cyclic-di-GMP signaling are required for polyamine-induced phage tolerance in *P. aeruginosa*.

**Figure S3.**
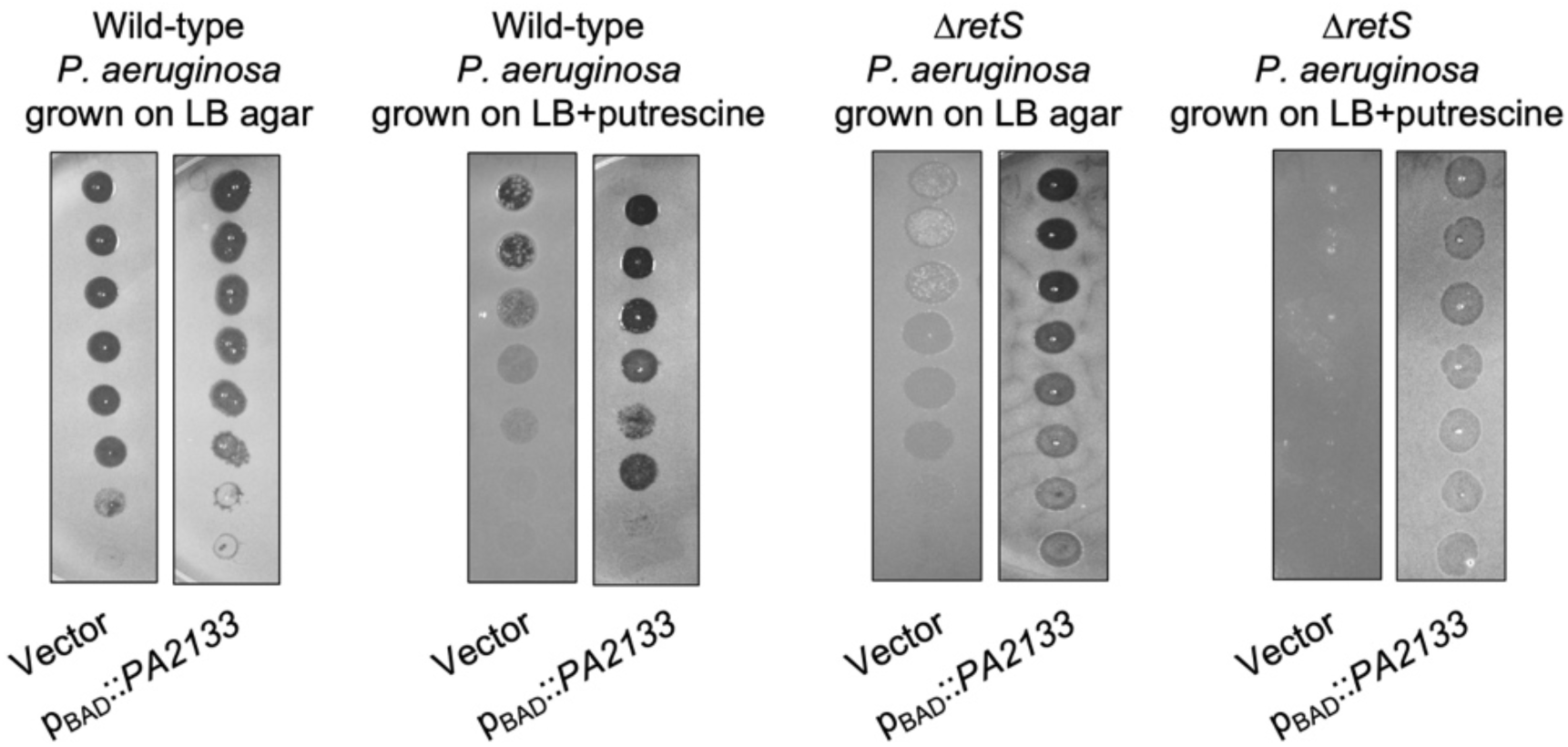
Expression of the cyclic-di-GMP degrading phosphodiesterase PA2133 restores sensitivity of *P. aeruginosa* to phage DMS3vir infection in the presence of putrescine. Serial dilution plaque assays comparing the plating efficiency of phage DMS3vir on the indicated *P. aeruginosa* strains carrying an inducible c-di-GMP-degrading phosphodiesterase (p_BAD_::*PA2133*) or a control strain containing an empty vector. Bacterial lawns were grown on LB agar supplemented with CHCl_3_-extracted lysate supplemented with 0.1% arabinose for 18 h.

### Putrescine suppresses DMS3vir genome replication and transcription

Intracellular polyamines are predominantly complexed with nucleic acids which affects many aspects of DNA replication and transcriptional regulation (Childs *et al*., 2003; Sarkar *et al*., 1995). We hypothesized that polyamines would interfere with phage DNA replication and/or transcription. To determine if polyamines affected phage DNA replication, we isolated DMS3vir DNA from infected cells with and without 50 mM putrescine after two hours and measured DMS3vir genome copy number by qPCR. In wild-type and Δ*retS P. aeruginosa*, putrescine inhibited DMS3vir genome replication by approximately 100-fold (**Fig 5A**). The effect was dependent on Gac/Rsm as DMS3vir genome replication was not affected in Δ*gacS* bacteria grown with putrescine compared to cultures grown without putrescine.

**Figure 5.**
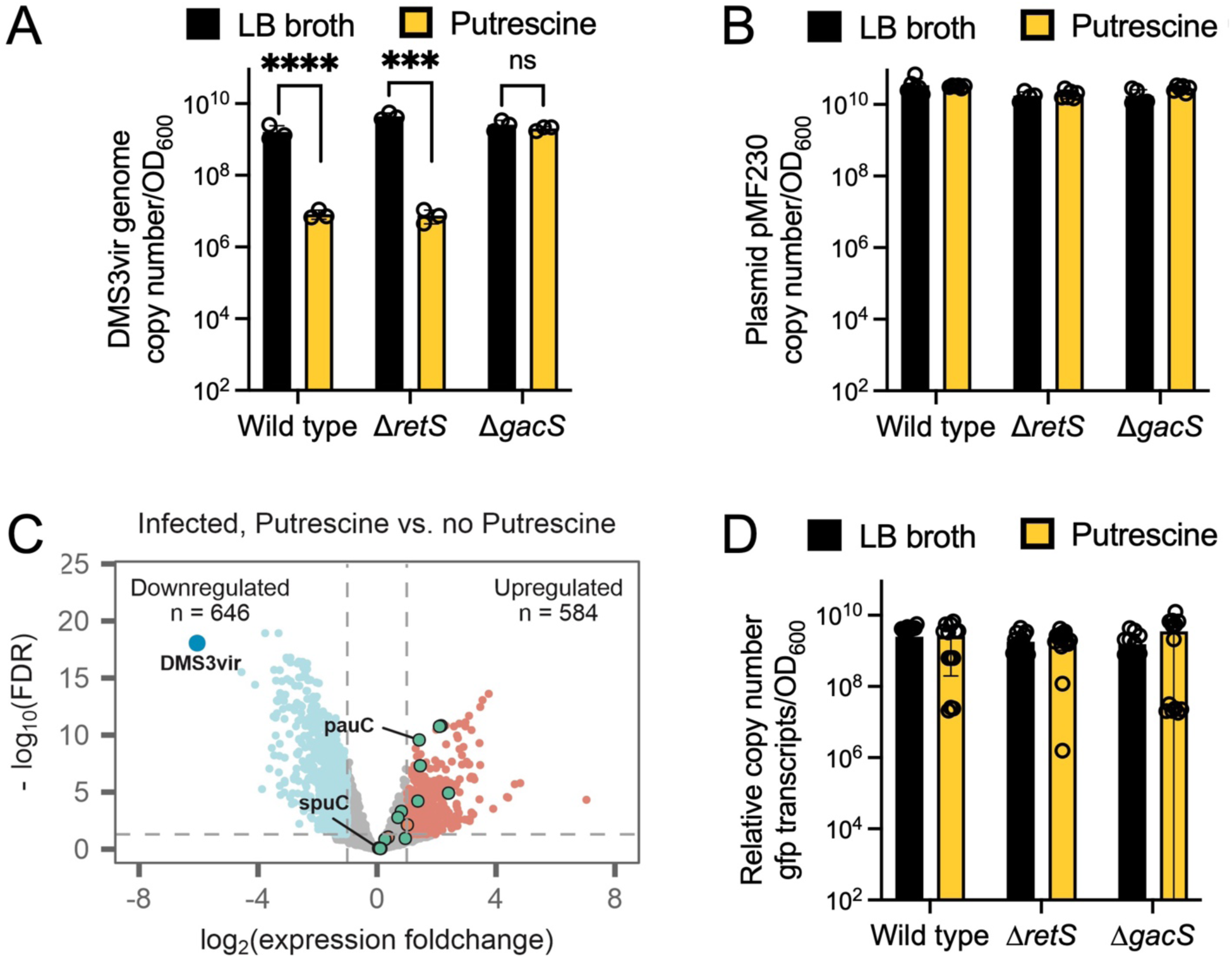
Putrescine inhibits DMS3vir genome replication and transcription but does not affect plasmid replication or transcription. **(A)** DMS3vir genome copy number was measured by qPCR in the indicated strains with or without 50 mM putrescine. Absolute copy number was determined using a standard curve generated with a cloned copy of the target sequence. DMS3vir copy number was then normalized to bacterial density (OD_600_). Results are mean ± SD of three experiments: ***P<0.001, ****P<0.0001, ns (not significant). **(B)** Plasmid pMF230 was purified from the indicated strains and copy number measured by qPCR (normalized to bacterial density OD_600_). Absolute copy number was determined using a standard curve generated with a cloned copy of the target sequence. Results are mean ± SD of six experiments: ***P<0.001. (**C)** Volcano plot showing differentially expressed genes in DMS3vir-infected *P. aeruginosa* treated with 50 mM putrescine vs non-treated infected cells. Red dots indicate upregulated genes (log_2_[foldchange] > 1 and FDR<0.05), and blue indicates downregulated genes (log_2_[fold change] < -1 and FDR<0.05). Non-significant genes are shown in gray. Genes involved in polyamine catabolism are highlighted. The large blue dot indicates reads that mapped to the DMS3vir genome. Data are representative of four experiments. (**D**) RT-qPCR was used to measure *gfp* expression in the indicated conditions and strains. Results are mean ± SD of at least 12 experiments. See also **Figure S4**.

Because the DMS3vir genome replicates as a circular episome, we hypothesized that polyamines would inhibit other episomally replicating DNA species such as plasmids. To test this, we measured the copy number of pMF230, a high copy number plasmid constitutively expressing GFP (Nivens et al., 2001). Putrescine did not affect plasmid copy number in wild-type, Δ*retS*, or Δ*gacS P. aeruginosa* (**Fig 5B**), disproving our hypothesis and suggesting that polyamines specifically inhibit phage but not plasmid DNA replication.

To test if polyamines affected phage transcription, we again used RNA-seq to measure the transcriptional response of *P. aeruginosa* infected by DMS3vir to exogenous putrescine. In DMS3vir-infected cells exposed to putrescine, several polyamine catabolism genes were upregulated and DMS3vir transcription was strongly downregulated compared to DMS3vir-infected cells not exposed to putrescine (**Fig 5C, Supplemental Table S1**). Conversely, transcription of *gfp* from pMF230 (as measured by RT-qPCR) was not significantly affected by putrescine in any strain or condition tested (**Fig 5D**), indicating that polyamines target phage transcription.

Collectively, these results indicate that in the presence of putrescine, DMS3vir successfully infects *P. aeruginosa*, initiates genome replication, but the DMS3vir lifecycle is disrupted at the level of DNA replication and transcription.

### Cyclic di-GMP genes are upregulated in DMS3vir-infected *P. aeruginosa* exposed to putrescine

Our data indicate that cyclic di-GMP signaling is required for polyamine-induced phage defense (**Fig 2D, Fig S3**). In *P. aeruginosa*, proteins that synthesize cyclic di-GMP contain a GGDEF domain whereas proteins with EAL domains are involved in cyclic di-GMP hydrolysis (Simm et al., 2004). We noted that eight genes encoding proteins with GGDEF and/or EAL domains were differentially regulated (seven upregulated and one downregulated) in phage-infected *P. aeruginosa* exposed to putrescine; phage infection by itself nor putrescine alone did not significantly affect the expression of genes encoding GGDEF and/or EAL domains (**Fig S4**). These results further implicate cyclic di-GMP signaling in mediating key biological processes related to polyamine-induced phage tolerance.

**Figure S4.**
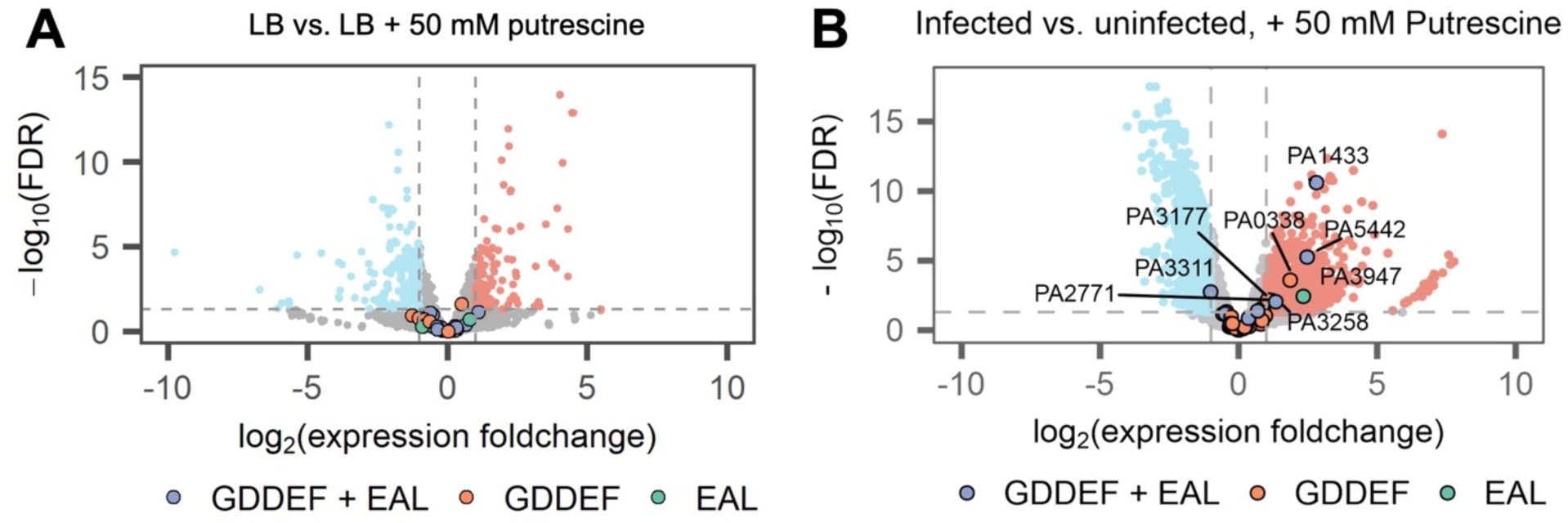
Phage infection in the presence of polyamines upregulates cyclic di-GMP regulation genes. Volcano plots showing differentially expressed genes in (**A**) *P. aeruginosa* cells grown in LB or LB supplemented with 50 mM putrescine or (**B**) DMS3vir-infected *P. aeruginosa* vs non-infected cells, both treated with 50 mM putrescine. Red dots indicate upregulated genes (log_2_[foldchange] >1 and FDR<0.05), and blue indicates downregulated genes (log_2_[fold change] <-1 and FDR<0.05). Non-significant genes are shown in gray. Genes involved in cyclic di-GMP metabolism are highlighted. Data are representative of four experiments.

### Phage infection induces Gac/Rsm-dependent intracellular polyamine accumulation

When transcriptional profiles from DMS3vir-infected and uninfected cells grown in the presence of putrescine were compared, we noted that polyamine catabolism genes were downregulated in infected cells, even though high levels of exogenous putrescine were present (**Fig S5**). We hypothesized that down regulation of polyamine catabolism genes in phage-infected cells would cause intracellular polyamine levels to increase. To test this, we measured total intracellular polyamine levels in pelleted and washed cells using a fluorometric assay. In uninfected wild-type, Δ*retS*, and Δ*gacS P. aeruginosa* growing in LB broth, basal intracellular polyamine levels were all approximately 8 mM after normalizing to bacterial density (OD_600_) (**Fig 6A–C**, black lines). In bacteria grown with 50 mM exogenous putrescine, intracellular polyamine levels spiked to over 40 mM/OD_600_ in wild-type and Δ*retS P. aeruginosa* during the first 30 minutes and returned to near basal levels over the course of 6 h (**Fig 6A and B**, green lines). Intracellular polyamine concentrations in the Δ*gacS* mutant did not increase in the first 30 minutes and remained at an elevated, albeit lower concentration (20 mM/OD_600_) over the course of the experiment (**Fig 6C**, green line).

**Figure 6.**
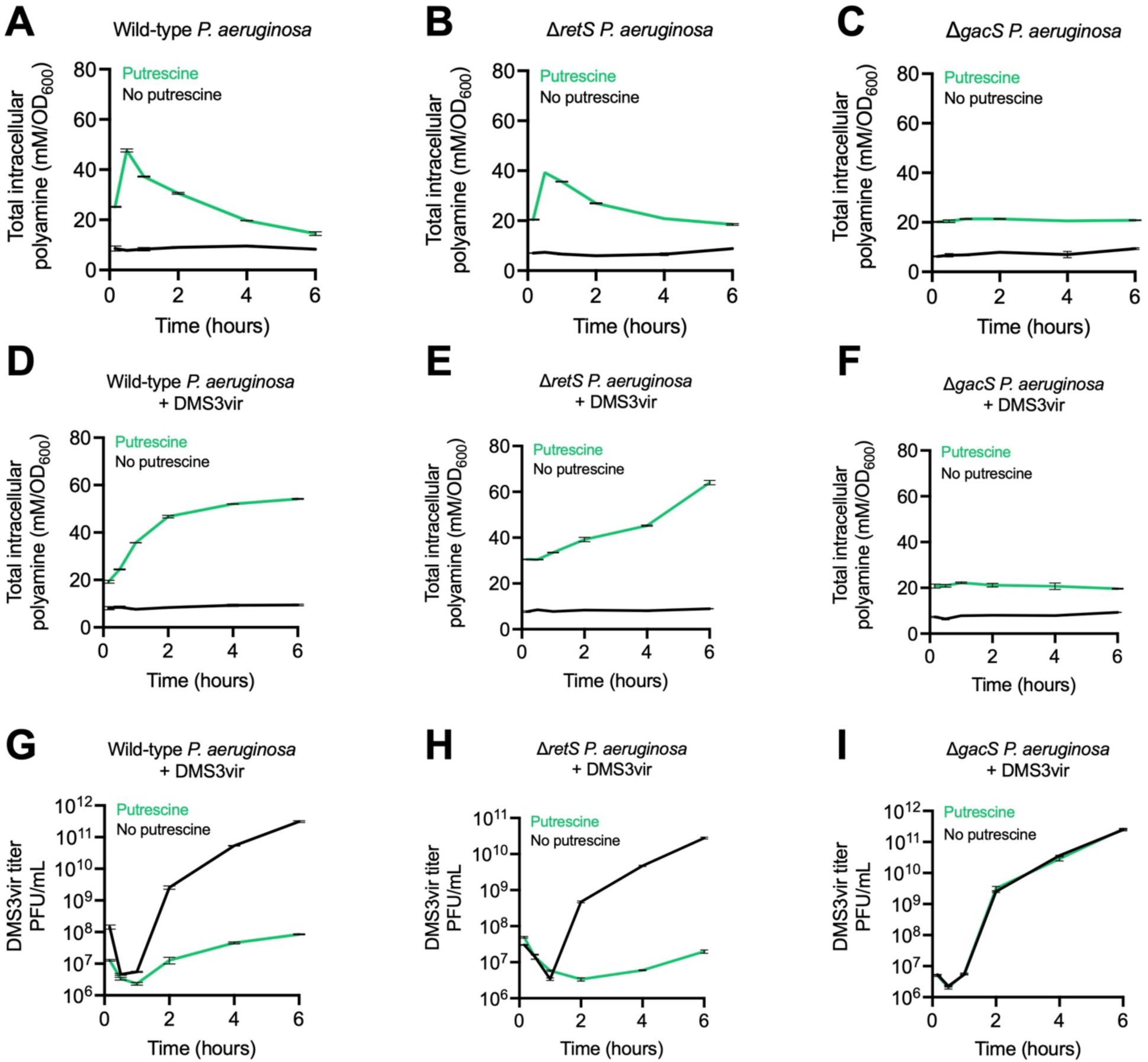
Gac/Rsm-dependent intracellular polyamine accumulation suppresses phage DMS3vir replication. Intracellular polyamines were measured in (**A-C**) uninfected cultures and (**D-F**) DMS3vir-innfected cultures in the indicated strains at the indicated times. Polyamine levels were normalized to bacterial density (OD_600_). Phage titers were also measured by plaque assay (**G–I**). Results are mean ± SD of duplicate experiments.

**Figure S5.**
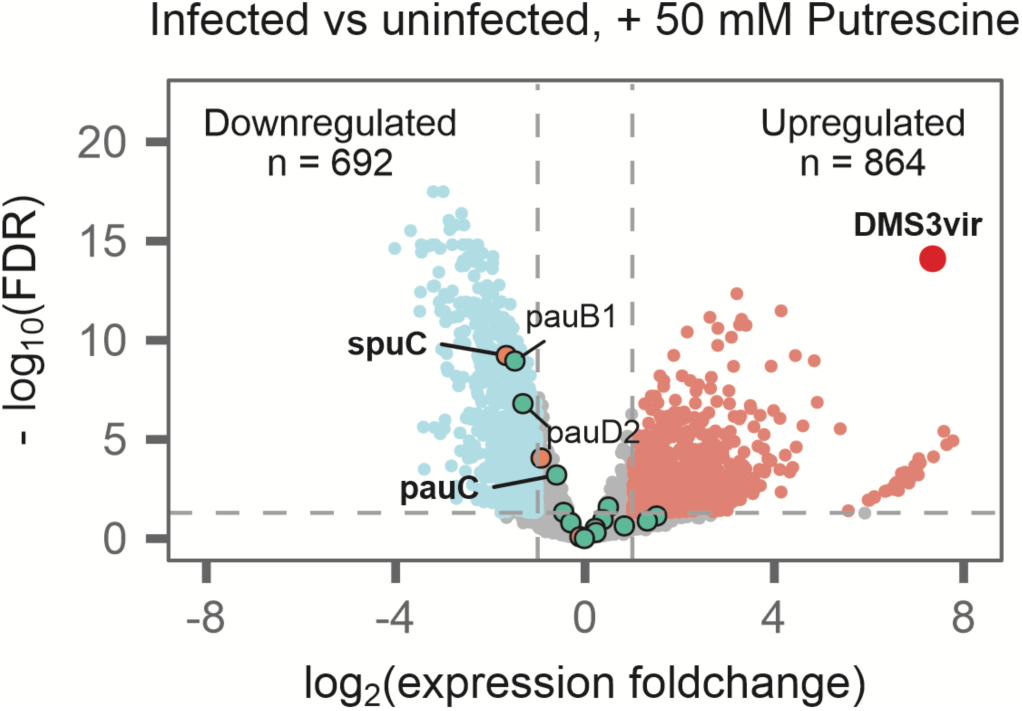
Volcano plot showing differentially expressed genes in DMS3vir-infected verses uninfected cells cultured with 50 mM putrescine. Red dots indicate upregulated genes (log_2_[foldchange] > 1 and FDR<0.05), and blue indicates downregulated genes (log_2_[fold change] < -1 and FDR<0.05). Non-significant genes are shown in gray. Genes involved in polyamine and putrescine catabolism are highlighted. The large red dot indicates reads that mapped to the DMS3vir genome. Four biological replicates for each condition are shown.

In response to infection by phage DMS3vir, intracellular polyamine levels remained at basal levels in wild-type, Δ*retS*, and Δ*gacS P. aeruginosa* growing in LB broth (**Fig 6D–F**, black lines). In the presence of exogenous putrescine, however, DMS3vir infection caused intracellular polyamine levels to increase to and remain at ∼50 mM/OD_600_ over the course of 6 h in both wild-type and Δ*retS P. aeruginosa* (**Fig 6D and E**, green lines). When Δ*gacS* was infected by DMS3vir in the presence of exogenous putrescine, intracellular polyamine levels remained constant at ∼20 mM/OD_600_ (**Fig 6F**, green line), comparable to uninfected Δ*gacS* cells shown in **Fig 6C**. This observation may explain why the Δ*gacS* strain has a polyamine-induced growth defect but is still sensitive to phage infection (see **Fig 4C**); intracellular polyamine levels may be sufficiently high to slow growth, but not high enough to inhibit phage replication.

We also measured intracellular polyamine levels in *P. aeruginosa* carrying plasmid pMF230. Because polyamines did not affect plasmid pMF230 replication or transcription (see **Fig 5B and D**), we predicted that this plasmid would not induce intracellular polyamine accumulation in *P. aeruginosa*. Indeed, in wild-type, *ΔretS*, and Δ*gacS P. aeruginosa*, pMF230 had no impact on intracellular polyamine accumulation (**Fig S6**).

Collectively, these results suggest that *P. aeruginosa* responds to the threat of phage infection by inducing Gac/Rsm-dependent intracellular polyamine accumulation.

**Figure S6.**
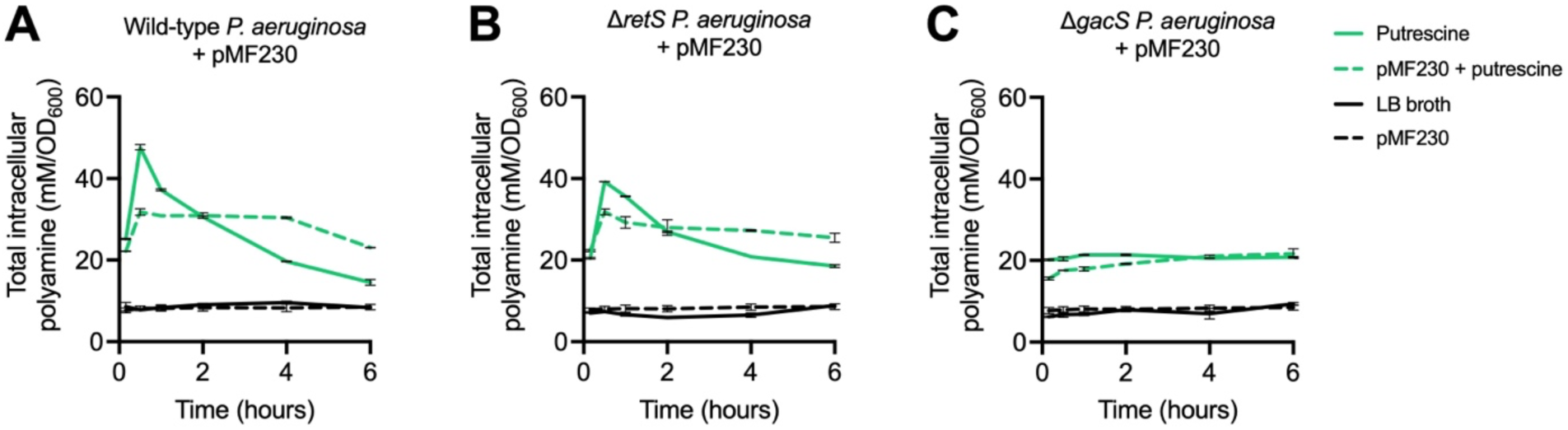
Putrescine does not inhibit plasmid pMF230 replication and pMF230 does not induce intracellular polyamine accumulation. (**A-C**) Intracellular polyamines were measured by fluorometric assay in the indicated strains grown in the presence or absence of 50 mM putrescine with or without the plasmid pMF230. Results are mean ± SD of duplicate experiments. Black lines represent cultures grown without putrescine; green lines represent culture supplemented with 50 mM exogenous putrescine. Solid lines represent uninfected cultures; dashed lines represent DMS3vir-infected cultures.

### Intracellular polyamine accumulation suppresses DMS3vir replication

In addition to intracellular polyamines, we also measured DMS3vir titers in bacterial supernatants collected from the cultures described above by plaque assay. In wild-type *P. aeruginosa* exposed to putrescine, DMS3vir titers were reduced by ∼3,500-fold compared to wild-type cells grown in LB broth after six hours (**Fig 6G**). Similar observations were made in Δ*retS P. aeruginosa*; in the presence of putrescine, DMS3vir replication in Δ*retS* was reduced ∼1,300 fold compared to Δ*retS* cultures without putrescine after six hours (**Fig 6H**). When Gac/Rsm signaling was disabled (Δ*gacS*), DMS3vir replication was not affected by putrescine and was comparable to DMS3vir replication in wild-type *P. aeruginosa* growing in LB broth (**Fig 6I**). These results indicate that Gac/Rsm-dependent intracellular polyamine accumulation suppresses DMS3vir replication.

### The N4-like phage CMS1 does not induce intracellular polyamine accumulation

Cell lysate and putrescine inhibited phages F8, DMS3vir, JBD26, and Pf4, but not phage CMS1 (**Fig 1A, Fig 3F and G**). CMS1 is a lytic N4-like phage in the family Podoviridae genus Litunavirus (Menon et al., 2021; Shi et al., 2020; Wagemans et al., 2014). Because CMS1 was able to replicate in a *P. aeruginosa* host in the presence of 50 mM putrescine, we reasoned that infection by CMS1 would not stimulate intracellular polyamine accumulation. Indeed, in contrast to DMS3vir, intracellular polyamine levels in CMS1-infected wild-type, Δ*retS*, or Δ*gacS P. aeruginosa* were comparable to uninfected cultures (**Fig 7A–C**; compare to **Fig 6A–C**). CMS1 produced a biphasic replication curve, similar to other N4-like phage species (Shi *et al*., 2020), and the presence of 50 mM exogenous putrescine had no significant impact on CMS1 titers compared to control cultures (**Fig 7D–F**). In addition to CMS1, the N4-like phage KPP21 (Shigehisa et al., 2016) was also not affected by exogenous putrescine (**Fig S7**). These results suggest that N4-like phages have evolved mechanisms to subvert intracellular polyamine accumulation phage defense.

**Figure 7.**
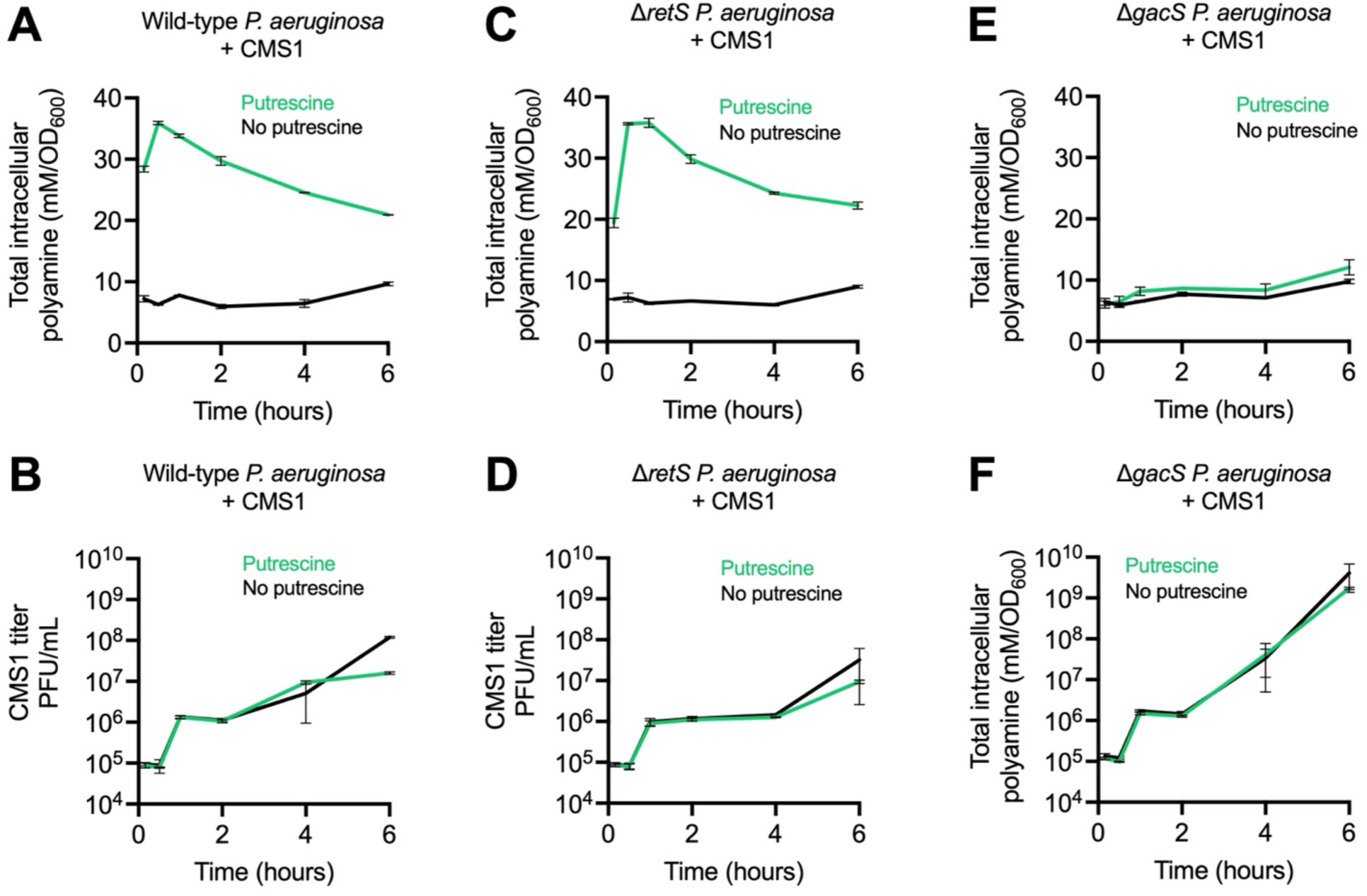
CMS1 infection suppresses Gac/Rsm-dependent intracellular polyamine accumulation. The indicated strains were grown to an OD_600_ of 0.3 in the presence or absence of 50 mM putrescine followed by infection with the N4-like phage CMS1 at a MOI of 1 where indicated. Intracellular polyamines and phage titers were then measured at the indicated times in **(A**,**B)** wild-type, **(C**,**D)** Δ*retS*, or **(E**,**F)** Δ*gacS P. aeruginosa*. Polyamine levels were normalized to bacterial density (OD_600_). Results are mean ± SD of duplicate experiments.

**Figure S7.**
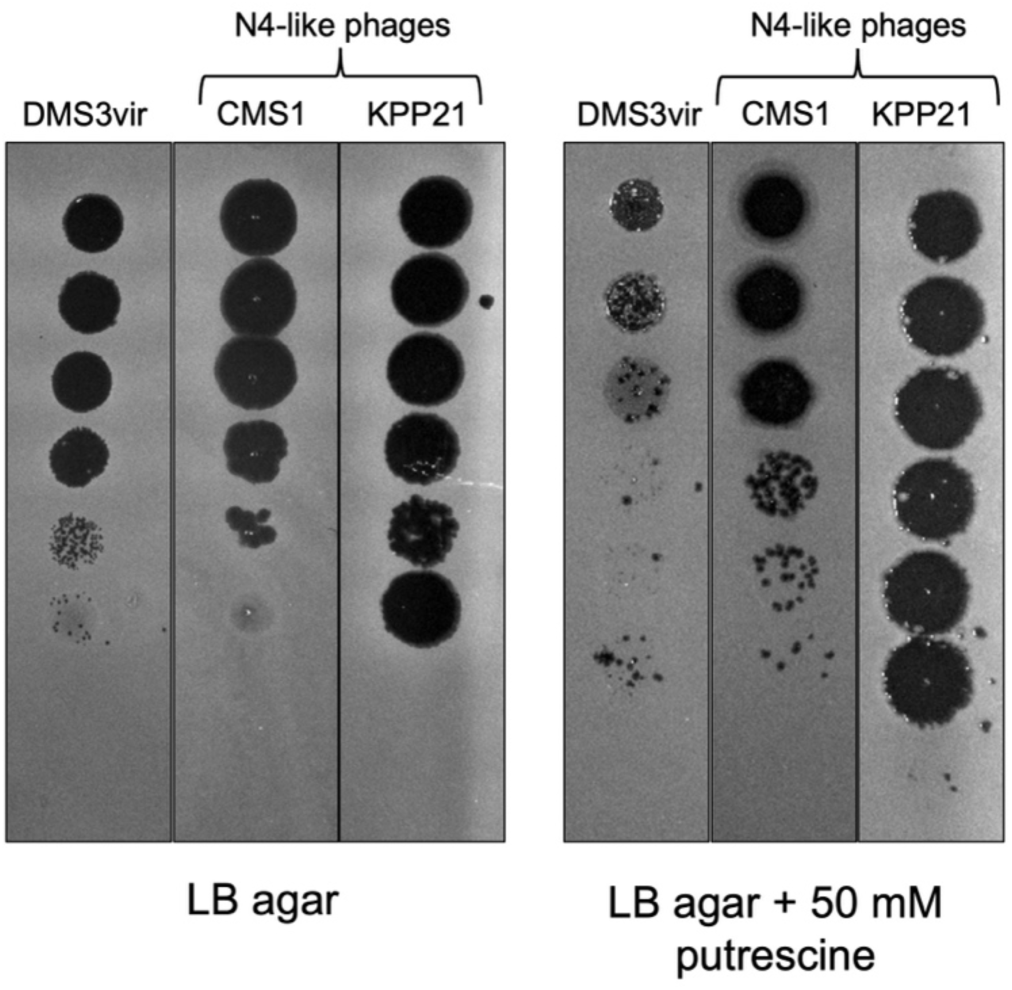
N4-like phages escape inhibition by exogenous putrescine. Serial dilution plaque assays comparing the plating efficiency of phage DMS3vir and the N4-like phages CMS1 and KPP21 on lawns of *P. aeruginosa* PAO1. Bacterial lawns were grown on LB agar or LB agar supplemented with 50 mM putrescine for 18 h.

## Discussion

In this study, we discovered a new phage defense system in which polyamines released by lysed bacteria serve as a damage signal that alerts *P. aeruginosa* of a nearby threat and prevents subsequent phage-mediated cell death. High exogenous polyamine levels result in bacterial intracellular polyamine accumulation. In the absence of a phage infection, intracellular polyamines are catabolized and return to basal levels. If *P. aeruginosa* is infected by a phage (i.e., a pathogen-associated signal is detected), polyamine catabolism genes are downregulated, intracellular polyamine levels remain elevated, and phage DNA replication and transcription are inhibited. Our data supports a model in which polyamines act as both a damage signal alerting nearby bacteria to cellular damage and as an inhibitor of phage replication.

Our results indicate a role for Gac/Rsm signaling in mediating phage defense in *P. aeruginosa*. The Gac/Rsm pathway in *P. aeruginosa* contains two major RNA binding proteins: RsmA and RsmN (Brencic and Lory, 2009; Romero *et al*., 2018). Prior work identified polyamine transport and metabolism genes in the RsmA regulon (the periplasmic polyamine-binding proteins PA2711 and PA0295 and the acetylpolyamine aminohydrolase PA1409) (Brencic and Lory, 2009). Polyamine catabolism genes are also in the RsmN regulon; the 5′-CANGGAYG motif recognized by RsmN is present in the polyamine metabolism genes *spuA, speC, pauB3, pauA5*, and *pauC* (Romero *et al*., 2018). Our results point to an unappreciated role for Gac/Rsm signaling in mediating intracellular polyamine homeostasis in *P. aeruginosa*.

Polyamines are ubiquitous throughout all domains of life and Gac/Rsm (CsrA) homologues are prevalent amongst γ-proteobacteria (Lapouge *et al*., 2008). Different organisms synthesize distinct types of polyamines and different polyamine species may induce different cellular responses in bacteria, which could provide context to bacterial threat assessments of cellular injury. Our work demonstrates that putrescine and spermidine (which are common in bacteria) induce cellular responses geared towards phage defense. In *Vibrio cholerae*, the periplasmic polyamine sensor MbaA can differentiate the relative abundance of the polyamines norspermidine and spermidine in the environment (Bridges and Bassler, 2021). Spermidine (but not norspermidine) activates MbaA signaling, which drives biofilm dispersal phenotypes. It is worth noting that *V. cholerae* MbaA periplasmic polyamine sensor contains both cyclic-di-GMP synthetase and phosphodiesterase domains that are essential for propagating the signal induced by endogenous spermidine (Cockerell et al., 2014). Because *V. cholerae* produces norspermidine but not spermidine, it is thought that norspermidine is perceived as a measure of “self” cell density while spermidine is perceived as a measure of how many “non-self” cells are present in the local environment (Sobe et al., 2017). The wide distribution of polyamines implicated in threat assessment suggests that these abundant intracellular metabolites may form the basis of a widely distributed threat assessment mechanism in bacteria.

Polyamines also influence bacterial responses related to virulence or immune evasion (Banerji *et al*., 2021). One example involves Gac/Rsm signaling and type III secretion gene regulation in *P. aeruginosa* (Mulcahy et al., 2006). The type III secretion apparatus delivers bacterial effectors to animal immune cells, allowing *P. aeruginosa* to escape phagocytic uptake (Yahr and Wolfgang, 2006). Exogenous polyamines such as spermidine and spermine (but not putrescine) induce type III secretion gene expression in *P. aeruginosa* (Williams McMackin et al., 2019; Zhou et al., 2007). Thus, different types of polyamines may guide bacterial responses to cellular injury caused by phagocytes or other components of the immune system.

Polyamines may serve as a damage signal in Gram-positive species as well. Recent work in *Bacillus subtilis* demonstrates that an uncharacterized soluble molecule released by lysed bacteria activates the stress-response sigma factor SigX, inducing the expression of enzymes that modify cell wall teichoic acids, conferring resistance to phage infection (Tzipilevich *et al*., 2021). It is possible that the low molecular weight soluble molecule that activates SigX in *B. subtilis* is a polyamine released by lysed kin cells.

Intracellular polyamine accumulation inhibited the replication of most but not all phages tested. The N4-like phages we tested were able to evade inhibition by polyamines by preventing intracellular polyamine accumulation. All known N4-like phages encode a cluster of small genes with mostly unknown function (Menon *et al*., 2021; Shigehisa *et al*., 2016). One of these genes in the N4-like phage LUZ7 genome encodes a protein called gp30 (Menon *et al*., 2021; Shigehisa *et al*., 2016). In a bacterial two-hybrid assay, gp30 directly interacted with the *P. aeruginosa* protein PA4114 (Wagemans *et al*., 2014), a spermidine acetyltransferase involved in polyamine catabolism. We speculate that modulating polyamine metabolism may be a common strategy used by N4-like phages to take over host cells.

How *P. aeruginosa* senses the threat of phage infection remains to be determined, but may involve the hostile takeover of essential bacterial complexes like RecBCD (Millman et al., 2020), alterations in bacterial respiration in response to phage infection (Carey et al., 2019), phage-induced membrane stress (Joly et al., 2010), or the detection of foreign nucleic acids (Datsenko et al., 2012). While foreign plasmid DNA does not induce intracellular polyamine accumulation (**Fig S4)**, it is possible that the linear dsDNA phage genome is detected as a threat as it is injected into a new host.

The ability to assess non-infectious and infectious threats is a fundamental feature of eukaryotic immune systems. In this study, we describe a new and more generalized way for *P. aeruginosa* to respond to infectious threats. Our study provides an additional example of how bacterial and eukaryotic immune systems are conceptually analogous in that they both sense and respond to damage- and pathogen-derived signals. Future studies to determine the precise mechanisms by which Gac/Rsm and cyclic-di-GMP signaling regulates intracellular polyamine accumulation and how bacterial cells sense phage infection will provide valuable new insights into phage-bacteria interactions that are relevant to many aspects of human health and disease.

## Methods

### Bacterial strains, plasmids, and growth conditions

Bacterial strains, plasmids, and their sources are listed in **Table 1**. Deletion mutants were constructed using allelic exchange and Gateway technology, as described previously (Fazli et al., 2015). Primer sequences are listed in **Table 2**. Unless indicated otherwise, bacteria were grown in lysogeny broth (LB) at 37°C with shaking and supplemented with antibiotics (Sigma) or 0.1% arabinose when appropriate. Unless otherwise noted, antibiotics were used at the following concentrations: gentamicin (10 or 30 µg ml^−1^), ampicillin (100 µg ml^−1^), and carbenicillin (50 µg ml^−1^).

**Table 1.** Bacterial strains, phage, and plasmids used in this study.

| Strain | Description | Source |
| --- | --- | --- |
| ***Escherichia coli*** |  |  |
| DH5α | Plasmid maintenance/propagation | New England Biolabs |
| S17 | λpir-positive strain used for conjugation | New England Biolabs |
| ***Pseudomonas aeruginosa*** |  |  |
| PAO1 | Wild type | (Schmidt et al., 2022) |
| PAO1 Δ*retS* | Clean deletion of *retS* from PAO1 | (Mougous et al., 2006) |
| PAO1 Δ*gacS* | Clean deletion of *gacS* from PAO1 | (Davies et al., 2007) |
| PAO1 Δ*rsmY/Z* | Clean deletion of *rsmY* and *rsmZ* from PAO1 | This study |
| PAO1 Δ*retS/rsmY/Z* | Clean deletion of *rsmY* and *rsmZ* from PAO1 Δ*retS* | This study |
| **Bacteriophage** |  |  |
| Pf4 | Inoviridae | (Secor et al., 2015) |
| JBD26 | Siphoviridae | (Bondy-Denomy et al., 2016) |
| CMS1 | Podoviridae | (Schmidt et al., 2022) |
| DMS3vir | Siphoviridae | (Cady et al., 2012) |
| F8 | Myoviridae | (Mendoza et al., 2020) |
| KPP21 | Podoviridae | (Shigehisa et al., 2016) |
| **Plasmids** |  |  |
| pJN105 | Empty expression vector with araC-P<sub>BAD</sub> promoter | (Hickman et al., 2005) |
| pJN105::PA2133 | PA2133 expression vector | (Hickman et al., 2005) |
| pMF230 | Constitutive expression of eGFP | (Nivens et al., 2001) |
| pEX18Gm | Allelic exchange suicide vector | (Hmelo et al., 2015) |

**Table 2.**
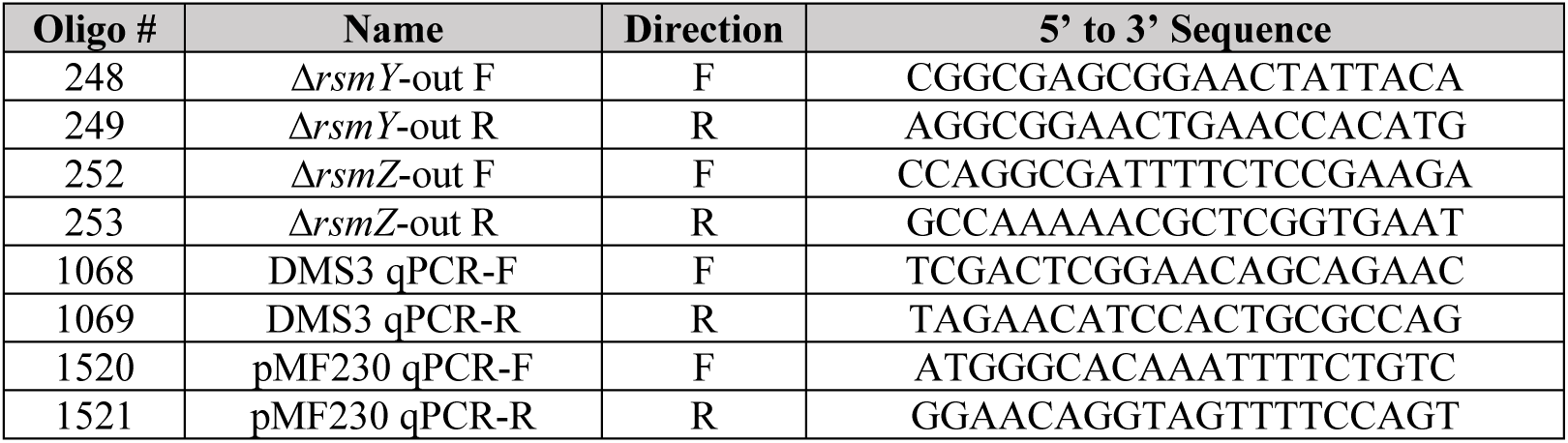
Primers used in this study.

### Cell lysate and polyamine agar plate preparation

*P. aeruginosa* PAO1 cells were pelleted, washed in PBS, and resuspended in fresh lysogeny broth (LB). Bacteria were then lysed by sonication and cell debris removed by centrifugation and filter sterilization. Agar plates were prepared by adding agar (1.5%) to the bacterial lysate. Polyamines (Putrescine dihydrochloride [MP Biomedicals] and Spermidine trihydrochloride [Sigma]) were added to sterile molten LB agar (1.5%) at the indicated final concentrations. Molten agar was mixed until polyamine powder was fully dissolved and then poured to make polyamine-supplemented agar plates.

### Plaque assays

Plaque assays were performed using lawns of the indicated strains grown on LB, cell lysate-supplemented, or polyamine-supplemented plates. Phages in filtered supernatants were serially diluted 10× in PBS and spotted onto lawns of the indicated strain. Plaques were imaged after 18 h of growth at 37°C.

### Growth curves

Overnight cultures were diluted to an OD600 of 0.05 in 96-well plates containing LB and, if necessary, the appropriate antibiotics, cell lysate, or polyamines. After 3 h of growth, strains were infected with indicated phage and growth measurements resumed. OD_600_ was measured using a CLARIOstar (BMG Labtech) plate reader at 37°C with shaking prior to each measurement.

### Polyamine measurements

Polyamines were measured using the Total Polyamine Assay Kit (MAK349, Sigma). A 100 µL aliquot of the indicated bacterial cultures was collected, centrifuged, washed with 1x PBS, and resuspended in PBS. Bacteria were lysed with 1:10 vol/vol chloroform, vortexed, and incubated at room temperature for 2 h. The solution was centrifuged, and the top aqueous layer was collected. 1.0 µL of the collected sample was mixed with the Total Polyamine Assay Kit reagents following the manufacturer’s instructions, incubated at 37°C for 30 minutes, and read using a CLARIOStar plate reader using end point fluorescence (λ_ex_ = 535 nm/λ_em_ = 587 nm). Polyamine concentrations were determined by comparing values to a standard curve constructed from known concentrations of putrescine. Values were then normalized to OD_600_ measurements taken from the original bacterial cultures.

### RNA purification and RNA-seq

Total RNA was extracted from the indicated strains and conditions using TRIzol. The integrity of the total cellular RNA was evaluated using RNA tape of Agilent TapeStation 2200 before library preparation. All RNA samples were of high integrity with a RIN score of 7.0 or more. rRNA was first depleted for each sample using MICROBExpress Kit (AM1905, Fisher) following the manufacture’s instruction. The rRNA-depleted total RNA was subjected to library preparation using NEBNext® Ultra™ II RNA Library Prep Kit (E7700, NEB) and barcoded with NEBNext Multiplex Oligos for Illumina (E7730, NEB) following the manufacturer’s instructions. The libraries were pooled with equal amount of moles, further sequenced using MiSeq Reagent V3 (MS-102-3003, Illumina) for pair-ended, 600bp reads, and de-multiplexed using the build-in bcl2fastq code in Illumina sequence analysis pipeline. Raw sequencing reads have been deposited as part of BioProject PRJNA806967 in the NCBI SRA database.

### RNA-seq data analysis

RNA-seq reads were aligned to the reference *P. aeruginosa* PAO1 genome (GenBank: GCA_000006765.1), mapped to genomic features, and counted using Rsubread package v1.28.1 (Liao et al., 2019). In viral challenge assays, the DMS3 phage genome (GenBank: DQ631426.1) was concatenated with PAO1 genome and treated as a single genomic feature. Count tables produced with Rsubread were normalized and tested for differential expression using edgeR v3.34.1 (Robinson et al., 2010) (**Supplemental Table S1**). Genes with ≥2-fold expression change and a false discovery rate (FDR) below 0.05 were considered significantly differential. Functional classification and Gene Ontology (GO) enrichment analysis were performed using PANTHER classification system (http://www.pantherdb.org/) (Mi et al., 2019). RNA-seq analysis results were plotted with ggplot2 and pheatmap packages in R.

### Fluorescence microscopy

*P. aeruginosa* strains of wild-type, Δ*gacS*, and Δ*retS* were back-diluted from overnight cultures and grown until OD_600_ = 0.2-0.3. Cells were then treated with putrescine to a final concentration of 50mM, DMS3vir, both, or neither. In the cultures that were treated with both, cells were pre-treated for 10 min with putrescine prior to the addition of phage. Cultures were then allowed to grow for an additional 2 h. Fluorescence microscopy was then performed as previously described (Brzozowski et al., 2019). Briefly, cells from culture aliquots of each strain in different treatment condition were stained with 1 µg/ml SynaptoRed (FM4-64) and 1 µg/ml DAPI to visualize the membrane and DNA respectively. Aliquots (5 µl) of the stained samples were then spotted onto glass bottom dishes (Mattek) and covered with a 1% agarose pad. Samples were imaged on a DeltaVision Elite microscope (Applied Precision/GE Healthcare/Leica Microsystems) equipped with a Photometrics CoolSnap HQ2 camera. Seventeen planes were acquired every 200 nm. The images were subsequently deconvolved using the manufacturer-supplied software, SoftWorx. Cell length was measured using ImageJ and analyzed in GraphPad Prism 9.

### Statistical analyses

Unless specified otherwise, differences between data sets were evaluated by Student’s *t* test, using GraphPad Prism version 5.0 (GraphPad Software, San Diego, CA). P values of < 0.05 were considered statistically significant.

## Acknowledgments

PRS was supported by NIH grants R01AI138981 and P20GM103546. PE was supported by NIH grant R35GM133617. BW was supported by NIH grant R35GM134867.

We declare no conflicts of interest.

